# Comparative genome analysis indicates rapid evolution of pathogenicity genes in *Colletotrichum tanaceti*

**DOI:** 10.1101/536516

**Authors:** Ruvini V. Lelwala, Pasi K. Korhonen, Neil D. Young, Jason B. Scott, Peter K. Ades, Robin B. Gasser, Paul W. J. Taylor

## Abstract

*Colletotrichum tanaceti* is an emerging foliar fungal pathogen of pyrethrum (*Tanacetum cinerariifolium*), posing a threat to the global pyrethrum industry. Despite being reported consistently from field surveys in Australia, the molecular basis of pathogenicity of *C. tanaceti* on pyrethrum is unknown. Herein, the genome of *C. tanaceti* (isolate BRIP57314) was assembled *de novo* and annotated using transcriptomic evidence. The inferred pathogenicity gene suite of *C. tanaceti* comprised a large array of genes encoding secreted effectors, proteases, CAZymes and secondary metabolites. Comparative analysis of its CAZyme pathogenicity profiles with those of closely related species suggested that *C. tanaceti* had additional hosts to pyrethrum. The genome of *C. tanaceti* had a high repeat content and repetitive elements were located significantly closer to genes inferred to influence pathogenicity than other genes. These repeats are likely to have accelerated mutational and transposition rates in the genome, resulting in a rapid evolution of certain CAZyme families in this species. The *C. tanaceti* genome consisted of a gene-sparse, A-T rich region facilitating a “two-speed” genome. Pathogenicity genes within this region were likely to have a higher evolutionary rate than the ‘core’ genome. This “two-speed” genome phenomenon in certain *Colletotrichum* spp. was hypothesized to have caused the clustering of species based on the pathogenicity genes, to deviate from taxonomy. With the large repertoire of pathogenicity factors that can potentially evolve rapidly in response to control measures, *C. tanaceti* may pose a high-risk to global pyrethrum production. Knowledge of the pathogenicity genes will facilitate future research in disease management of *C. tanaceti* and other *Colletotrichum* spp..

## INTRODUCTION

Plant pathogens cause diseases world-wide that have devastating economic, social and ecological consequences [1]. Fungi are among the dominant causal agents of plant diseases [2] and the genus *Colletotrichum* has been ranked among the top-ten most important fungal plant pathogens [3]. Many *Colletotrichum* species are known to cause major economic losses globally, and have been extensively used in the study of the molecular and cellular bases of fungal pathogenicity [4]. The publication of 25 whole genome sequences of *Colletotrichum* species has significantly improved understanding of the biology, genetics and evolution of this genus [5–11]. However, a large research gap still exists with this ever-expanding genus consisting of more than 200 accepted species [12] and 14 major species complexes [13, 14]. The availability of only one genome of a member of the destructivum complex, *C. higginsianum*, [5, 15] has constrained comparative studies within and among species complexes. Insights into the genomic organization and the pathogenicity gene repertoire of other *Colletotrichum* species in the destructivum complex therefore, will significantly expand the knowledge base of this important genus.

*Colletotrichum tanaceti*, a member of the destructivum complex [16], is an emerging foliar fungal pathogen [17] of Dalmatian pyrethrum *(Tanacetum cinerariifolium)*. Pyrethrum is commercially cultivated as a source of the natural insecticide pyrethrin [18]. *Colletotrichum tanaceti* has been consistently reported in Australian field surveys of the crop [19] since 2013 [17] and causes leaf anthracnose, with black, water-soaked, sunken lesions [17]. Due to its hemibiotrophic lifestyle, characteristic symptoms of *C. tanaceti* are not evident on leaves until around 120 hours after infection [17, 20], when it switches from biotrophy to necrotrophy. A significant reduction in green leaf area occurs usually 10 days after infection [17]. This suggests a rapid disease cycle for *C. tanaceti* in pyrethrum and, given its aggressiveness, the potential for serious crop damage.

The molecular basis of pathogenicity of *C. tanaceti*, which includes the pathogenicity genes and their evolution, has not been studied. *Colletotrichum tanaceti* has only been reported from pyrethrum in Australia, but may have crossed over from another plant host species. However, cross-host pathogenicity has not yet been assessed and the pathogen’s origin and the potential host range are currently unknown. Therefore, the threat posed by *C. tanaceti* to the local and global pyrethrum industry remains largely unknown.

The genome sequence of an emerging plant pathogen such as *C. tanaceti* can provide a foundation for identifying genes associated with the pathogen life cycle, pathogenicity and virulence. Effectors [21], proteases [22], and carbohydrate active enzymes (CAZymes) that [23] are important gene categories in fungal pathogenesis. Furthermore secondary metabolites and transporters, *P450s* and transcription factors [24] associated with biosynthesis of secondary metabolites are also important pathogenicity factors. Fungal mitogen activated protein (MAP) kinase pathways regulate the cascade of reactions that respond to various environmental stresses and are also important factors determining pathogenicity and virulence [25]. Draft genomes of many fungal pathogens have been used to infer genes involved in pathogenicity with a high accuracy [26, 27] using homology searches against curated databases [28, 29] and *de novo* inference using bioinformatics tools [21, 30]. Therefore, characterization of the genome of *C. tanaceti*, followed by inference and quantification of these important pathogenicity gene categories will be beneficial for future functional and pathogenicity studies of this and related pathogens.

Comparative genomics has enabled inference of patterns of speciation, pathogenesis and host determination within *Colletotrichum* lineages [31]. These studies have indicated that the gain and loss of putative pathogenicity gene families in *Colletotrichum* genomes are important determinants of host specificity and pathogenic adaptation of these species [7, 11].

Comparison of putative pathogenicity gene repertoires of *Colletotrichum* species from different species complexes and species closely related to the genus *Colletotrichum* would provide insights into the evolutionary rates of these genes. Comparative genomics will also enable the quantification of pathogenicity at the molecular level and identification of the host range of *C. tanaceti* with respect to other *Colletotrichum* species. Therefore, combined genomics and comparative genomics analyses can provide sound means of assessing the current and future risks posed by *C. tanaceti*.

In order to achieve the major goal of evaluating the potential threat to the pyrethrum industry form *C. tanaceti*, the aims of this study were to: 1) infer the pathogenicity gene suite of *C. tanaceti*; 2) quantify the molecular basis of pathogenicity; 3) infer the host range of *C. tanaceti*; and 4) quantify the rate of evolution of pathogenicity genes in *C. tanaceti*.

## MATERIALS AND METHODS

### Sequencing and *de novo*-assembly of the genome of *C. tanaceti*

#### Fungal strain

The ex-holotype of *C. tanaceti* strain BRIP57314 (CBS 132693=UM01) [17] was acquired from the culture collection of BRIP (Plant Pathology Herbarium, Department of Primary Industries, Queensland, Australia). This isolate was propagated on potato dextrose agar (PDA; Sigma Aldrich, St. Louis, USA) and incubated at 24°C using a 12 h:12 h light:dark photoperiod. Genomic DNA was isolated using a modified CTAB protocol [32]. The integrity and quantity of DNA was confirmed by 1.5% agarose gel electrophoresis and a nanodrop spectrophotometer (Thermo Fisher Scientific, Waltham, USA)

#### Genome sequencing and assembly

Genomic DNA was fragmented using a Covaris ultrasonicator (Covaris Inc., Massachusetts, USA) to achieve an average fragment length of 532 base pairs (bp). A genomic DNA library with an average insert size of 420 bp was constructed using the KAPA Hyper Prep Library Preparation Kit [33] and was paired-end sequenced (2×300 bp reads) using the Illumina Miseq platform (San Diego, USA). The raw reads were filtered for low quality nucleotides and adapters using Trimmomatic [34] (Phred score-33, leading-3, trailing-6, slidingwindow-4:15, minlen-36) to retain 22,871,341 sequences and were profiled using KAT [35]. Filtered reads were then assembled using DISCOVAR *de novo* [36]. The completeness of the assembly was assessed with the Sordaromyceta_*odb9* gene set [37] using the program Benchmarking Universal Single-Copy Orthologs (BUSCO v2) [37] in the Genomics Virtual Laboratory platform [38]. The GC-bias of the genome was detected using OcculterCut version 1.1 with default settings [39].

#### Prediction of repetitive elements

Species-specific repeats were first inferred using the program RepeatModeler [40], in which the programs RECON [41] and RepeatScout [42] were used. Long terminal repeats (LTRs) were predicted using the program LTR_Finder [43]. The program RepeatMasker v4.0.5 [44] was employed to mask resulting species-specific repeats and LTRs; and applied the program Tandem Repeat Finder (TRF) [45] and the database Repbase v.17.02 [46] to predict and mask interspersed and simple repeats. All repeats predicted were combined using ProcessRepeats command in RepeatMasker.

### RNA sequencing

#### Inoculation of Pyrethrum leaves

Pyrethrum leaves were inoculated using the leaf-sandwich method [47, 48] by placing a fungal ‘mat’ between two pyrethrum leaves in a petridish. Each petri dish was sealed with parafilm and incubated at 24°C with a 12 h-photoperiod. Induced mycelia were harvested at 6, 24 and 48 h after inoculation, and total RNA was extracted using the RNeasy Plant Mini kit (Qiagen, Australia) following the manufacturer’s instructions. Total RNA was extracted from the saprobic stage (1-week-old cultures growing on potato dextrose agar). Contaminating genomic DNA was removed from RNA samples by Ambion™ DNase I (Thermo Fisher Scientific, USA) treatment; the integrity and quantity of total RNA was confirmed by 1% agarose gel electrophoresis and the Experion^™^ automated electrophoresis system (Biorad Laboratories, Australia).

RNA libaries were prepared using both E7530L and E&335L NEBNext^®^ Ultra^™^ RNA Library Prep Kits (New England Biolabs, USA) to generate fragment sizes of 351-371 bp. The transcriptome was paired-end sequenced (2 × 150 bp reads) on the Illumina Hiseq 2500 platform (San Diego, USA). Raw reads were trimmed for quality using Trimmomatic [34] (leading-25, trailing-25, slidingwindow-4:25, minlen-40) to retain between 17,935,938 – 18,761,773 sequences for each library and profiled using FastQC [49].

#### Gene prediction

Genes were first predicted using the MAKER3 v3.0.0-beta [50], in which both the transcriptomic data from *C. tanaceti* and the proteomic and *ab initio* gene predictions from *C. graminicola;* [51] and *C. higginsianum;* [51] were combined into a consensus prediction. In brief, transcriptomic RNAseq reads of *C. tanaceti* were assembled into transcripts in both *de novo* and genome-guided modes of the program Trinity v2.2.0 [52]. In genome guided assembly, reads were mapped onto the genome using the program TopHat2 v2.1.0 [53]. Genome guided and *de novo* transcriptomic assemblies were combined, redundancy (99% similarity) was removed using the program cd-hit-est [54, 55] and resulting transcripts were filtered for full-length open reading frames (ORFs) using the program Transdecoder [52]. Resulting full-length transcripts were further reduced to 80% similarity using the program cd-hit-est and checked for splicing sites. These high quality transcripts were then used as a training set for *ab initio* gene prediction programs AUGUSTUS v3.1 [56] and SNAP v6.7 [57] and GENEMARK v4.2.9 [58]. Evidence data from assembled transcriptomes (with 99% redundancy using cd-hit-est) and the proteomes were provided to Maker3. The predicted genes (length of conceptually translated protein ≥ 30 amino acids) were further clustered using the *k*-means clustering algorithm [59] with following metrics: 1) Maker3 annotation edit distance (AED); 2) number of exons in the mRNA; 3) length of translated protein sequence; 4) fraction of exons that overlap transcript alignment; 5) fraction of exons that overlap transcript and protein alignment; 6) fraction of splice sites confirmed by a SNAP prediction from Maker3; 7) percentage for repeat overlap with gene-, exon- and CDS-sequence; 8) size of the inferred orthologous group the gene belongs to using OrthoMclv2.0.9 [60]; and 9) presence of functional annotation (see Functional annotation of the *C. tanaceti* genome section below). Resulting clusters with transposons and *ab initio* gene predictions with no transcriptome or proteome support were removed.

#### Functional annotation of the *C. tanaceti* genome

Putative coding regions were subjected to protein homology searches against the NCBI (nr) and Swiss-Prot database using BLAST v 2.7.1 (E-value of ≤ 1e-8) [61]. Conserved protein domains and gene ontology (GO) terms were assigned to predicted proteins using InterProScan 5 [62]. Additionally, Kyoto Encyclopedia of Genes and Genomes (KEGG) Orthology (KO) terms were assigned to predicted proteins using the Blastkoala search engine [63]. Assigned KO terms were used to generate *C. tanaceti* pathway maps using KEGG mapper [64]. Putative genes of *C. tanaceti* with functional annotations were subjected to species-specific gene enrichment analysis on the DAVID functional annotation database tool [65, 66] and using *C. graminicola* as the reference species.

#### Comparison to related taxa

The genome and proteome of *C. tanaceti* was compared to genomes of related taxa using genome alignment, synteny and orthology analyses as following.

#### Genome alignment and synteny analysis

*Colletotrichum tanaceti* genome contigs were aligned to 13 other publicly available genomes (Table 1) of *Colletotrichum* species using nucmer in Mummer v 3.9.4 [67]. Contig-alignments were then filtered for a minimum 30% nucleotide identity and 200 bp in aligned length. The global coverage of each of the genomes by contigs of *C. tanaceti* was computed. The program ‘Synteny Mapping and Analysis Program’, SyMAP v 4.2 [68] was used to map *C. tanaceti* contigs (>150 kb) to the genome with the highest coverage to identify the syntenic regions which are the regions that are in preserved order in chromosomes of *C. higginsianum* IMI349063 reference genome [5].

**Table 1.** Genomes used in the comparative genomic analyses

| Organism | Identifier<sup>a</sup> | Taxonomy ID<sup>b</sup> | Genbank accession number<sup>c</sup> | Bio project ID<sup>d</sup> | Strain<sup>e</sup> | Assembly<sup>f</sup> | Reference |
| --- | --- | --- | --- | --- | --- | --- | --- |
| *Colletotrichum chlorophyti* | CCh | 708187 | MPGH000000000.1 | PRJNA350752 | NTL11 | ASM193710v1 | [69] |
| *Colletotrichum fioriniae* | CFi | 1445577 | JARH000000000.1 | PRJNA233987 | PJ7 | GCA_000582985.1 | [70] |
| *Colletotrichum fructicola* | CFr | 1213859 | ANPB000000000.1 | PRJNA225509 | Nara gc5 | GCA_000319635.1 | [6] |
| *Colletotrichum gloeosporioides* | CGl | 1237896 | AMYD000000000.1 | PRJNA176412 | Cg-14 | GCA_00446055.1 | [71] |
| *Colletotrichum graminicola* | CGr | 645133 | ACOD000000000.1 | PRJNA37879 | M1.001 | M1_0001_v1 | [5] |
| *Colletotrichum higginsianum* | CHi | 759273 | LTAN000000000.1 | PRJNA47061 | IMI349063 | GCA_001672515.1 | [5] |
| *Colletotrichum incanum* | CIn | 1573173 | LFIW000000000.1 | PRJNA286717 | MAFF238704 | GCA_001189835.1 | [9] |
| *Colletotrichum nymphaeae* | CNy | 1460502 | JEMN000000000.1 | PRJNA237763 | IMI504889 | GCA_001563115.1 | [7] |
| *Colletotrichum orchidophilum* | COc | 1209926 | MJBS000000000.1 | PRJNA411788 | IMI309357 | GCF_001831195.1 | [72] |
| *Colletotrichum orbiculare* | COr | 1213857 | AMCV000000000.1 | PRJNA171217 | MAFF240422 | Corbicular240422v01 | [6] |
| *Colletotrichum salicis* | CSa | 1209931 | JFFI000000000.1 | PRJNA238477 | CBS607.94 | GCA_001563125.1 | [7] |
| *Colletotrichum*<br>*simmondsii* | CSi | 703756 | JFBX0000<br>0000.1 | PRJNA<br>239224 | CBS<br>122122 | GCA_001563135 | [7] |
| *Colletotrichum*<br>*sublineola* | CSu | 1173701 | JMSE0000<br>0000.1 | PRJNA<br>246670 | TX430<br>BB | GCA_000696135.1 | [73] |
| *Colletotrichum*<br>*tanacetii* | CT1 | 1306861 | PJEX0000<br>0000 | PRJNA<br>421029 | BRIP57<br>314 |  |  |
| *Verticillium*<br>*dahliae* | VDh | 498257 | ABJE0000<br>0000.1 | PRJNA<br>225532 | VdLs.1<br>7 | GCF_000150675.1 | [74] |
| *Botrytis cinerea* | BCi | 332648 | AAID000<br>00000.2 | PRJNA<br>15632 | B05.10 | GCF_000143535.2 | [75] |
| *Sordaria*<br>*macrospora* | SMa | 771870 | CABT000<br>00000.2 | PRJNA<br>51569 | k-hell | GCF_000182805.2 | [76] |
| *Fusarium*<br>*oxysporum* | FOx | 426428 | AAXH000<br>00000.1 | PRJNA<br>18813 | CBS<br>123668 | GCF_00149955.1 | [77] |

<sup>a</sup> Short identifier used in place of the species name in supplementary information
<sup>b</sup> Taxonomy ID of each species according to the NCBI taxonomy database
<sup>c</sup> Genbank accession number for the deposited nucleotide sequence
<sup>d</sup> NCBI bioproject ID
<sup>e</sup> version of the genome assembly

#### Orthology search and phylogenomics analysis

The proteomes of *C. tanaceti* and the publicly available 17 other species (Table 1) were subjected to ortholog searching using OrthoMCL v2.0.9 [60] and MCL [78] with an inflation value of 1.5. The orthoMCL output was used to determine the percent orthology among the species and to determine the core gene set for *Colletotrichum*. The ortho-groups with pathogenicity genes (inferred as below) of *C. tanaceti* were extracted and used to determine the percent conservation of those gene categories within the genus. Furthermore the single copy orthologs were extracted from the orthoMCL output and aligned using MAFFT v.7 [79]. These alignments were then trimmed using trimAl v.1.3 [80] to remove all positions in the alignment with gaps in 20% or more of the sequences, unless this leaves less than 60% of the sequence remaining. The trimmed reads were concatenated using FASconCAT-G [81]. The concatenated alignment was partitioned and amino acid substitution models were predicted for each partition using ProtTest 3 [82] in FASconCAT-G. The partitioned, concatenated alignment was subjected to maximum likelihood phylogenetic analysis using RAxML v8.2.10 [83] to find the best tree from 20 maximum likelihood searches and using 100 bootstrap replicates. Evolutionary distance in number of substitutions per site was computed using the *ape* package [84] in the R statistical language framework v 3.5.1. [85] from the maximum likelihood tree.

#### Estimation of divergence dates

The phylogram developed from above was utilized to estimate the divergence dates of the species considered as following. The final RAxML phylogenetic tree was used to generate an ultrametric tree in r8s v1.81 [86] applying the penalized likelihood method [87] and the truncated Newton (TN) algorithm [88]. Divergence times were estimated using previously derived estimates [8, 11, 89] of 267-430 million years (Myr) for the Leotiomycetes-Sordaromycetes crown, 207-339 Myr for the Sordaromycete crown and 45-75 Myr for the *Colletotrichum* crown as calibrations. An optimal smoothing factor which was deduced using the cross validation process [86] among 50 values across 1 to 6.3e+09 was used in the divergence time estimation.

#### Prediction of secretome and database searches for identifying other virulence factors

Predicted proteins of *C. tanaceti* were used in downstream prediction of the secretome [90]. A combination of three software tools: SignalPv4.1 [91], Phobius [92] and WoLFPSORT [93] was used to predict the signal peptides. Proteins with transmembrane domains were identified using TMHMM v.2.0 [94] and were excluded as secreted proteins. Proteins with signals targeting the endoplasmic reticulum and GPI anchors were identified and excluded using Ps-SCAN [95] and Pred-GPI [96] respectively. NLStradamus [97] was used to identify proteins with nuclear localization signals. Curated secretome was subjected to homology search against the CDD database to identify the conserved domains (E-value ≤ 1e-10). The candidate secreted effector proteins were identified by passing the secretome through the program EffectorP [21]. Predicted effector candidates were manually inspected and candidates with known plant cell wall degrading catalytic domains, such as cutinases (PF01083.21), short-chain dehydrogenases (PF00106.24), glycosyl hydrolases (PF00457), peptidases (PF04117.11) and lipases (PF13472.5) were excluded. The candidates with no detectable conserved domains and no homology (E-value ≤ 1e-3) to any other proteins in NCBI– non-redundant protein sequence database were defined as species-specific. Putative secreted peptidases and inhibitors were predicted by stand-alone blastp (E-value ≤ 1e-10) homology searches of the domain database of MEROPS release 12.0 [98]. Furthermore, potential virulence factors of *C. tanaceti* were identified by blastp searches (E-value ≤ 1e-10) against PHI-base v 4.4 [28]. The online analysis tools, Antibiotics and Secondary Metabolite Analysis Shell (antiSMASHV.4) [30] with default parameters and SMURF [99] were used to predict potential secondary metabolite backbone genes and clusters using the default parameters. Cytochrome P450s and transporters were described based on blastp (E-value ≤ 1e-10) homology searches against the Fungal Cytochrome P450 database [100] and the Transported Classification Database [101]. The functional annotations for *C. tanaceti* were compared across 17 other closely related taxa (Table1). The family specific Hidden Markov Model profiles of dbCan database v6 [102] were employed using the program HMMScan in HMMER v31.b2 [103] in order to identify the carbohydrate active enzymes (CAZymes) and the CAZyme families in the proteome of *C. tanaceti*. Fungi specific cut-off E-value of 1e-17 and a coverage cut-off of 0.45 [102] were used in the analysis which was repeated for seventeen related species (Table 1). The identified CAZymes were run though InterProScan 5 [62] to check for false positives. The member counts of each CAZyme family for each taxon were corrected accordingly.

#### Evolution of CAZyme gene families

CAFE v4.0 [104, 105] was used to estimate the number of CAZyme gene family expansions, contractions and the number of rapidly evolving gene families upon divergence of different lineages. Error-models [105] were estimated to account for the genome assembly errors and were incorporated into computations. A universal lambda value (maximum likelihood value of the birth-death parameter) was assumed and gene families with significant size variance were identified using a probability value cut-off of 0.01. The branches responsible for significant evolution, were further identified using the Viterbi algorithm [104] with a probability value cutoff of 0.05. Sizes of plant pathogenicity-related gene families from CAZomes of each of the species; the ‘CAZyme pathogenicity profiles’ were retrieved and compared using the online tool ClustVis [106]. The ‘CAZyme pathogenicity profile’ of a particular species included the gene families that have activities in binding to or degradation of plant cell wall components such as cellulose, hemicelluloses, lignin, pectin and cutin.

#### Relationship of pathogenicity related genes and repeat elements

The mean distances between repetitive elements and pathogenicity related genes were analyzed using permutation tests implemented in the package regioneR [107] in the R. Repetitive element categories incorporated in this analysis included: 1) tandem and interspersed repeats combined; 2) tandem repeats; and 3) interspersed repeats. These were compared to the pathogenicity related gene classes: 1) CAZymes; 2) peptidases; 3) secondary metabolite biosynthetic gene clusters; and 4) effectors. The mean distance between each gene in above categories and the nearest repetitive element was compared against a distribution of distances of random samples from the whole genome. Ten thousand random iterations were conducted, from which a Z-statistic estimate, and its associated probability, were computed for each gene category.

## RESULTS

### *Colletotrichum tanaceti* genome and gene content

The genome of isolate BRIP57314 was assembled into 5,242 contigs with an N50 value of 103,135 bp and assembly size of 57.91Mb. The average GC content was 49.3% (Table 2). The genome size and GC content of *C. tanaceti* was within the range previously reported to other *Colletotrichum* spp. (S1 Fig). Draft genome assembly and the raw unassembled sequences are available under the accession no PJEX00000000 in Genbank. The genome contained 12,172 coding genes with an average gene length of 2,575bp. Mean exon count per gene was 3, and 54.1% of the genome sequence contained protein-encoding genes. In the BUSCO analysis, out of the 3,725 benchmarking genes in the Sordaromyceta group, the genome was reported to contain 3,656 complete BUSCOs (98.2%), of which two were duplicated and the rest were single copy genes (98.1%). A total of 30 (0.8%) BUSCOs were fragmented and 39 were missing (1.0%). The repeat content of *C. tanaceti* was 24.6% of the total genome of which 85.2% was interspersed repeats (Table 3).

**Table 1.** Features of the *Colletotrichum tanaceti* BRIP57314 genome

| Feature | Statistics |
| --- | --- |
| GC_content (%) | 49.3 |
| N50 (bp) | 103,135 |
| Maximum sequence length (bp) | 945,015 |
| Mean length (bp) | 11,047 |
| Number of base pairs | 57,912,474 |
| Number of contigs | 5,242 |
| Number of genes | 12,172 |
| Number of exons | 35,792 |
| Number of introns | 23,620 |
| Number of CDS | 12,172 |
| Overlapping genes | 3,983 |
| Contained genes | 1,586 |
| Mean gene length (bp) | 2,575 |
| Mean exon length (bp) | 787 |
| Mean intron length (bp) | 137 |
| Mean CDS length (bp) | 1,440 |
| % of genome covered by genes | 54.1 |
| % of genome covered by CDS | 30.3 |
| Mean mRNAs per gene | 1 |
| Mean exons per mRNA | 3 |
| Mean introns per mRNA | 2 |

**Table 3.** Repetitive elements of the *C. tanaceti* genome

| Repetitive element | Number of elements | Length occupied (bp) | Percentage of sequence |
| --- | --- | --- | --- |
| SINEs: | 49 | 4,123 | 0.01 |
| ALUs | 0 | 0 | 0 |
| MIRs | 11 | 869 | 0 |
| LINEs: | 612 | 251,619 | 0.43 |
| LINE1 | 207 | 48,554 | 0.08 |
| LINE2 | 35 | 2,588 | 0 |
| L3/CR1 | 82 | 5,928 | 0.01 |
| LTR elements: | 7,299 | 4,825,086 | 8.33 |
| ERVL | 2 | 120 | 0 |
| ERVL-MaLRs | 1 | 39 | 0 |
| ERV_classI | 3 | 209 | 0 |
| ERV_classII | 1 | 32 | 0 |
| DNA elements: | 1,436 | 905,846 | 1.56 |
| hAT-Charlie | 3 | 140 | 0 |
| TcMar-Tigger | 6 | 529 | 0 |
| Unclassified: | 8,863 | 6,153,241 | 10.62 |
| Total interspersed repeats |  | 12,139,915 | 20.96 |
| Small RNA: | 754 | 210,370 | 0.36 |
| Satellites: | 0 | 0 | 0 |
| Simple repeats: | 9,941 | 1,757,918 | 3.04 |
| Low complexity: | 3,064 | 147,883 | 0.26 |
| Total repeat content |  |  | 24 .62% |

Of the 12,172 predicted proteins, 11,352 had an annotation edit distance (AED) value of less than 1.0, and 2962 genes had an AED value of zero. The number of genes without putative annotation from the public database searches was only 958. A total of 8,945 proteins (73.5% of proteome) had InterProScan annotations of which 6,911 contained 9,647 Pfam domain annotations and 5,452 had GO term ontology annotation. The most abundant (*n*=129) Pfam domain was the cytochrome P450 family (PF00067) followed by the protein kinase domain (*n*=127; PF00069). Gene enrichment analysis suggested enrichment of many GO terms including those associated with translation and chromosome telomeric region (S1 Table). Putative proteins of *C. tanaceti* were subjected to KEGG pathway analysis which returned assignment of 5,883 proteins to known pathways (S2 Table). The highest number of KO identifiers was among the metabolic pathway assignments (*n*=693) of which the majority (*n*=363) were for amino acid metabolism followed by carbohydrate metabolism (*n*=290) (S3 Table). Among the environmental information processing pathways, 81 *C. tanaceti* genes were assigned into 47 KO identifiers belonging to MAPK pathway (S4 Table). Furthermore, 24 *C. tanaceti* proteins were annotated with 10 aflatoxin biosynthesis pathway KO assignments (S5 Table) and 56 proteins were assigned KOs for ABC transporters (S6 Table).

### Genome alignment and synteny

The global alignment coverage of 13 other *Colletotrichum* genomes from *C. tanaceti* contigs was proportionate to the evolutionary proximity to *C. tanaceti* (Fig 1a). The highest coverage was in *C. higginsianum* (63.8%) and the least was in *C. orbiculare* 4.26%. Among the *C. tanaceti* contigs aligned to the chromosomes of *C. higginsianum*, the best alignment coverage was to chromosome NC_030961.1 (chromosome 9) (S7 Table). *Colletotrichum tanaceti* contigs (*n*=155 of size≥10 kb) were mapped in SyMAP synteny analysis to form 142 synteny blocks which covered 44.0% of the *C. higginsianum* and 80.0% of the *C. tanaceti* sequences that were used (S2 Fig). Genes were present in 92.0% of the syntenic regions in *C. tanaceti* and in 77.0% of *C. higginsianum*. No inverted synteny blocks were reported. Despite the highest coverage in *C. higginsianum* chromosome 9, the largest synteny block was identified between the complete *C. tanaceti* contig 4 (945.01 kb of length) and *C. higginsianum* chromosome NC_030954 (Chromosome 1). A total of 38 effector candidates of *C. tanaceti* were within these syntenic regions between *C. tanaceti* and *C. higginsianum*. No synteny blocks were detected to the two mini chromosomes (NC_030963.1 and NC_030964.1) of *C. higginsianum*.

**Fig 1.**
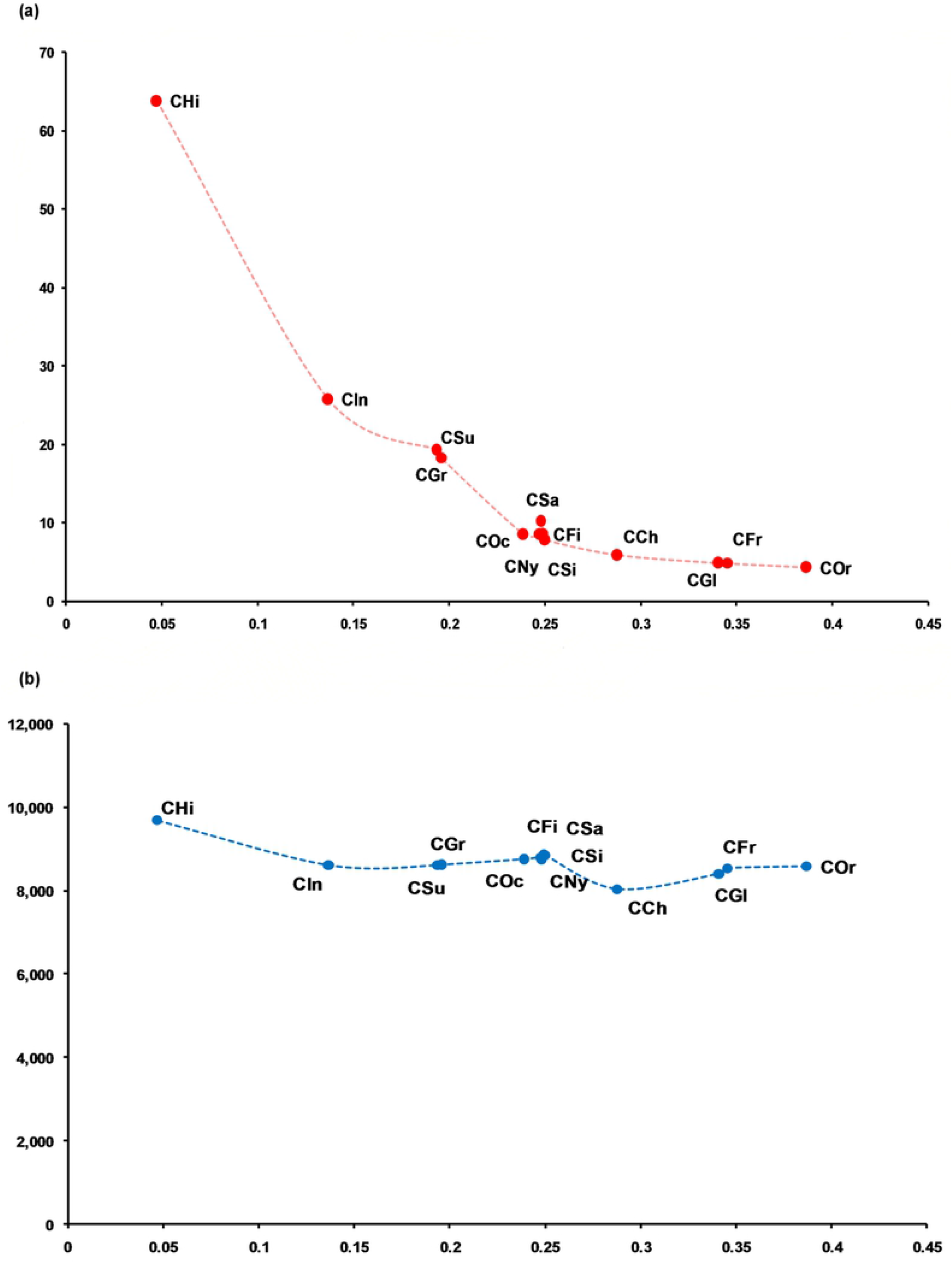
Comparison of the *C. tanaceti* genome to previously published *Colletotrichum* spp. genomes. (a) Percentage global alignment (y axis) of 13 *Colletotrichum* draft genomes to contigs representing the *C. tanaceti* draft genome, plotted against evolutionary distance with reference to *C. tanaceti* (x axis), (b) Number of orthologs shared by 13 *Colletotrichum* draft genomes and *C. tanaceti* (y axis) plotted against the evolutionary distance with reference to *C. tanaceti* (x axis); evolutionary distance given in number of substitutions per site, computed using the ape package [98] in R from a maximum likelihood tree.

### Orthology search

Of 221,456 total genes from 18 genomes, the number of core genes reported for all ascomycetes in the orthology analysis was 3,944. A total of 10,695 putative proteins from *C. tanaceti* were assigned to 10,074 groups containing orthologs and/or recent paralogs and/or co-orthologs across all species. A total of 6,002 genes were conserved in the genus *Colletotrichum. Colletotrichum tanaceti* had 9,679 orthologs with *C. higginsianum* which was the highest ortholog count among *Colletotrichum* spp. followed by 8,855 orthologs with *C. nymphaea* (Fig 1b). Twenty of these groups, with 48 genes among them were exclusive to *C. tanaceti* and were defined as recent paralogs (*in-paralogs*) of *C. tanaceti* with no homology to the 16 other species tested.

### Divergence time in *Colletotrichum* lineages

A total of 2,214 single copy ortholog (SCO) genes identified among the *C. tanaceti* and 17 closely related genomes (Table 1) were used to generate a maximum likelihood (ML) evolutionary tree in which all branches achieved bootstrap support of 100%. *Colletotrichum tanaceti* formed a clade with *C. higginsianum*, a member of the destructivum complex and the two destructivum complex members formed a sister clade with the graminicola complex members and *C. incanum*. A smoothing factor value of 1 was reported as the optimal value for divergence time predictions in r8s. *Colletotrichum tanaceti* and *C. higginsianum* were reported to have diverged ~ 9.97 million years ago (mya). The most recent common ancestor (MRCA) of gloeosporioides, graminicola, and acutatum clades were reported to be 6.12, 10.98 and 15.78 mya, respectively (Fig 2).

**Fig 2.**
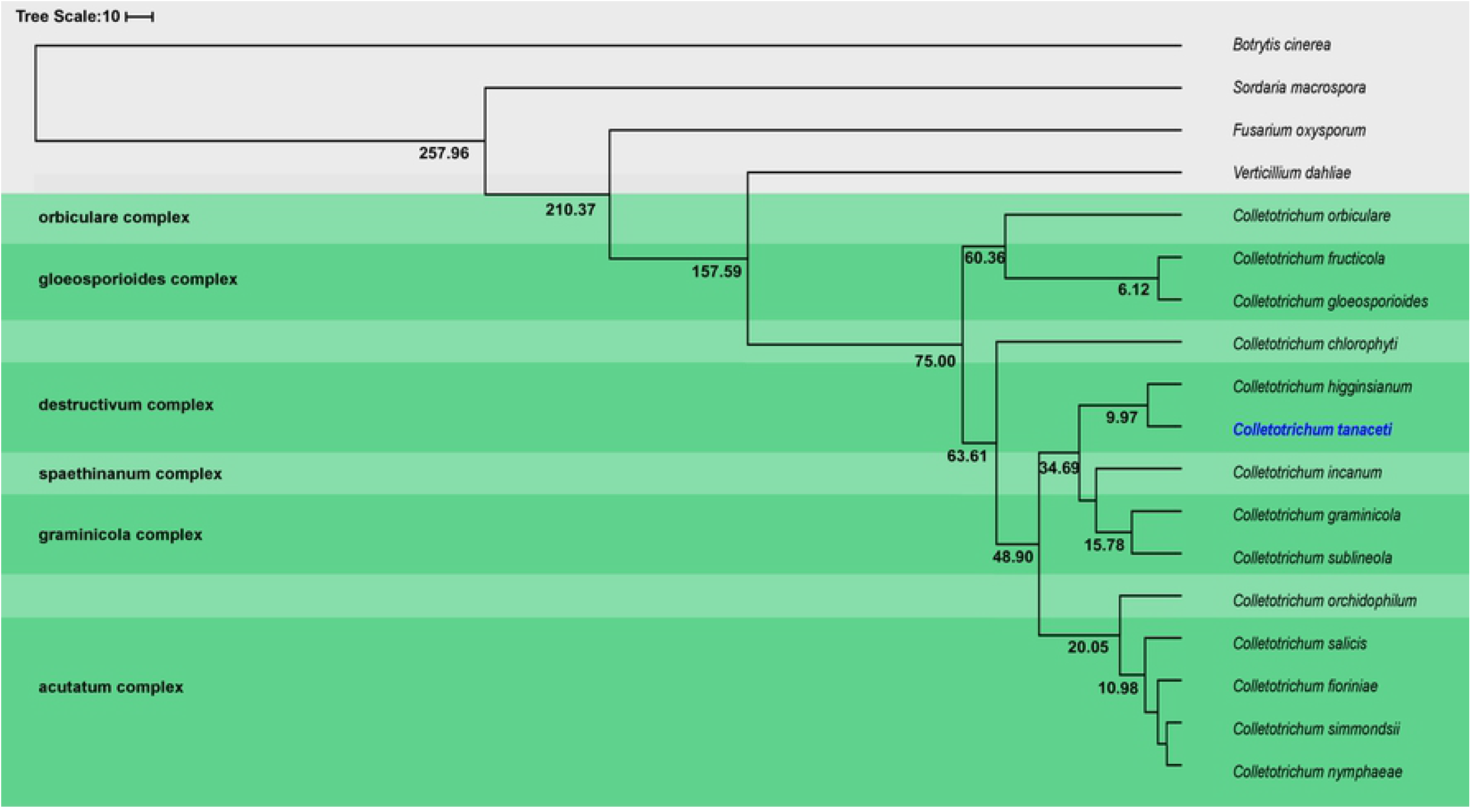
Chronogram showing divergence time estimations (in million years) for *Colletotrichum* spp. and related taxa.

### Identification of pathogenicity related genes in *C. tanaceti*

#### Secretome of *C. tanaceti*

Of the 12,172 predicted proteins, 1,024 (8.41%) were predicted to be secreted. A total of 2,702 Conserved Domain Database (CDD) domains were found in the secretome. Of these, 287 were specific features with NCBI curated models, 124 were generic features with only the superfamily annotations [108]. Only 433 queries had no known domain hits. The secretome was rich in alpha beta hydrolase superfamily (cl21494) containing enzymes, glycosyl hydrolases and proteolytic enzymes and cytochrome P450 monoxygenases (*P450*) (S8 Table). A total of 100 secreted proteins had nuclear-localization signals (S8 Table).

A total of 233 effector candidates were predicted by EffectorP. Following manual inspection and censoring for candidates with known plant cell wall degrading catalytic domains, a total of 168 candidates were selected as *C. tanaceti* effector candidates for further analysis (S9 Table). The secreted candidate effector repertoire of *C. tanaceti* contained homologs of known effectors, such as the *Ecp6* of *Cladosporium fulvum* [109], *MC69* of *Magnoporthe oryzae* [110], *ToxB* of *Pyrenophora tritici repentis* [111] and *Magnoporthe oryzae Bas3* [112]. Furthermore, among the effector candidates, there were proteins with conserved domains of known virulence factors. Most effector candidates were small (average length of 155 amino acids) and rich in cysteine (average cysteine composition was 3.3%) which are the hallmarks of effectors. A total of 78 conserved motifs of fungal effectors [113] were present in 62 effector candidates which had at least one motif each. Twenty-two effector candidates that did not cluster in ortholog search among the 14 *Colletotrichum* and three related species, and also did not show detectable homology to the NCBI-nr and swissprot databases were defined as *C. tanaceti*-specific. Only 24% of the effectors of *C. tanaceti* were conserved among all 14 *Colletotrichum* spp.

A total of 98 secreted peptidases were predicted with the majority (*n*=64) being serine peptidases largely comprising the S08 and S09 subfamilies. The second most abundant class was the metallo peptidases (*n*=19) (S10 Table). All six aspartic peptidases belonged to subfamily A01. A total of 20 secreted peptidase inhibitors were reported in *C. tanaceti* comprising two carboxypeptide-y inhibitors, five family-19 inhibitors and 13 family-14 inhibitors (S11 Table). Forty nine percent of the proteases of *C. tanaceti* were among the “core” set of proteases of *Colletotrichum*.

#### Secondary metabolite-related genes and clusters

Forty-one secondary metabolite backbone genes were predicted in *C. tanaceti* using SMURF and the majority were polyketide synthases (PKs, *n*=13) with four PKs-like proteins. Furthermore, nine non-ribosomal peptide synthases (NRPS), eight NRPs–like proteins, two hybrid PKs-NRPS enzymes and five dimethylallyltryptophans (DMATS) were also predicted as backbone genes (S12 Table). A total of 52% of these backbone genes were within the core set of genes in *Colletotrichum*. A total of 33 secondary metabolite gene-clusters were predicted surrounding the backbone genes. However, the program antiSMASH predicted a total of 50 clusters. Among the clusters, there were 12 type1-PKS, two type3-PKs, thirteen terpenes, eleven NRPS, four indoles, three T1pks-nrps, one T1PKs-indole and four other proteins. Cluster 10 of T1PKS showed 100% similarity to the genes in LL-Z1272 beta biosynthetic gene cluster (BGC0001390_c1). Furthermore, a homolog to the melanin biosynthetic gene *SCD1* was also reported in *C. tanaceti* (CTA1_6632). When predictions from the two tools were compared, putative SMB clusters on 31 contigs of *C. tanaceti* were predicted by both tools and 19 of the backbone genes from SMURF were also predicted in antiSMASH (Supplementary Table 12). A total of 37 SM clusters were within the syntenic blocks of *C. higginsianum*. The conserved SM domains identified in each cluster were reported (S13 Table). Predictions from antiSMASH were compared across taxa and majority of the clusters were type1-PKs like followed by NRPS in all ascomycetes compared (Fig 3a). The highest number of clusters were reported from *C. fructicola* (*n*=84) followed by *C. higginsianum* (*n*=74) and *C. gloeosporioides* (*n*=73). The composition of the SMB gene cluster composition of *C. tanaceti* was most similar to *C. orchidophilum*, the acutatum complex members and *C. orbiculare* (S3 Fig).

**Fig 3.**
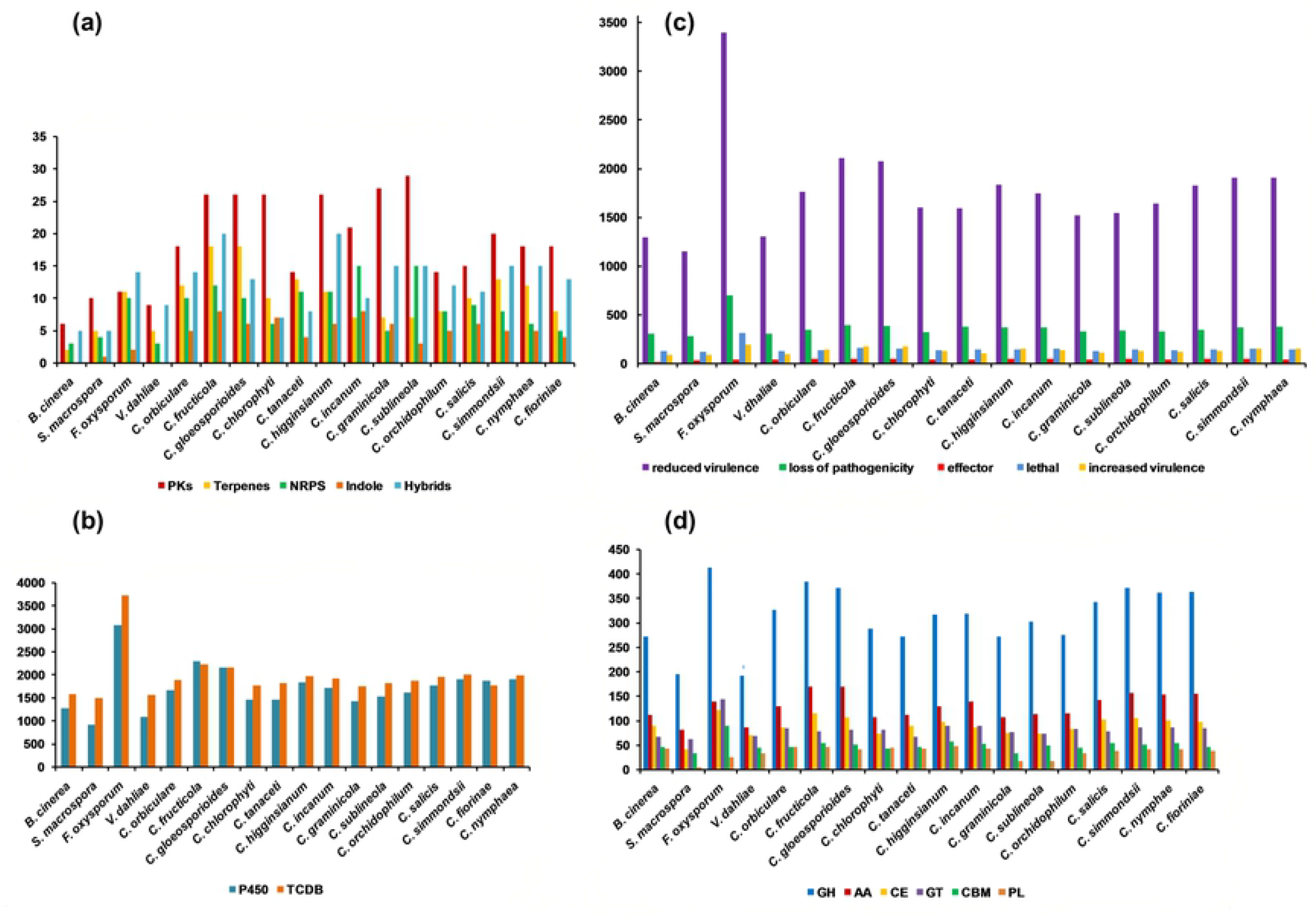
Composition of different pathogenicity gene categories predicted for *Colletotrichum tanaceti* and related species. The number of genes in each gene category (x axis) plotted for each species (y axis). (a) secondary metabolite biosynthetic gene clusters- (gene clusters producing polyketides, terpenes, non-ribosomal peptides (NRPs), indoles and the hybrids of above); (b) number of homologs in the fungal cytochrome P450 database and the transporter classification database (TCDB); (c) homologs in the pathogen-host interaction database; homologs to entries in the “unaffected pathogenicity” database were excluded; (d) CAZyme classes; glycoside hydrolases (GH), polysaccharide lyases (PL), glycosyltransferases (GT), carbohydrate esterases (CE), molecules with auxiliary activities (AA), and carbohydrate binding molecules (CBM).

#### Cytochrome P450 monoxygenases (P450s) and transporters

In the *C. tanaceti* genome, 1,457genes had homologs in the fungal cytochrome P450 database (S14 Table) and 911 out of that had >30% identity. There were 1,824 homologs (S15 Table) in the transport classification database for *C. tanaceti* with 1,276 genes with >30% identity. The majority (*n*=430) of the homologs were genes of the major facilitator superfamily (MFS, 2. A.1) followed by 129 genes of the ABC transporter family (3.A.1) and 123 of N.P.C 1.I.1. Within *Colletotrichum* genus, members of the gloeosporioides complex had the highest number of homologs for both P450s and transporters (Fig 3b).

#### Homologs in PHI-base

A total of 3,497 homologs were recorded in *C. tanaceti* from the pathogen-host interaction database (PHI), of which 1,592 represented mutated phenotypes with reduced virulence (S16 Table). The second most common (*n*=1,514) were the unaffected pathogenicity category, 382 homologs were for loss of pathogenicity and 42 were in the effector category. Notably, the mutant phenotype of 141 homologs was lethal to this particular pathogen, and 103 homologs had increased virulence after mutation (Fig 3c). The two gloeosporioides complex members had the highest number of homologs in the database among the *Colletotrichum* spp., followed by the acutatum complex species, *C. simmondsii, C. fioriniae* and *C. nymphaea*. Despite *C. higginsianum* having a large number of homologs, *C. tanaceti* had a below average number for all the categories among the *Colletotrichum* spp., with a profile similar to *C. orchidophilum*, *C. chlorophyti* and *C. graminicola* (S4 Fig).

#### CAZymes

A total of 608 *C. tanaceti* proteins were assigned to 121 CAZyme families of which 43% was glycosyl hydrolases followed by 18 % of redox enzymes (auxiliary activities) and 14% carbohydrate esterases (S17 Table). Carbohydrate binding molecules and polysaccharide lyases both formed 7% each of the *C. tanaceti* CAZome whereas 11% was glycosyltransferases. Members of the gloeosporioides and acutatum complexes had the largest CAZomes among *Colletotrichum* spp. The CAZyme repertoires of the graminicola complex members were relatively small (Fig 3d).

#### Evolution of CAZyme families upon divergence of *Colletotrichum* lineages

A total of 152 CAZyme families, predicted at the node of MRCA for *S. macrospora* and *B. cinerea*, were used in gene family evolution analyses in CAFÉ. A uniform birth-death parameter (*λ*) of 0.0023 was computed. Thirty gene families were reported to be significantly evolving (family-wide *p* value ≥ 0.05), of which 21 were rapidly evolving (family-wide *p*≥ 0.01 and *Viterbi p* ≥ 0.01 in any lineage) (S18 Table).

At the divergence of *Colletotrichum* spp., 39 expansions and 12 contractions were predicted with respect to its MRCA with *Verticillium* species (S19 Table). Expansions included the lignin hydrolase family AA2, pectin degrading polysaccharide lyase families (PL1, 3, 4, 9 and GH78), lignocellulose degrading families (AA3, AA9, GH131, GH5, GH6, GH7), hemicelluloses degrading families (CE1, CE4, CE5, CE12, GH3, GH16, GH30, GH43, GH51, GH67, and GH10), Lys M domain containing family CBM50 and cutinase family CE5. The cellulose degrading family GH131 was the only rapidly evolving CAZyme family (family-wide *p* ≥ 0.01 and *Viterbi p* ≥ 0.01) which expanded upon the divergence of *Colletotrichum* spp. Within the genus, the highest number of expansions (*n*=38) was reported at the divergence of the gloeosporioides-complex clade with only 4 contractions. Notably, the CBM18 and GH10 families were contracted and many families with plant cell wall degrading enzyme activity were expanded. The rapidly and significantly expanded families, (family-wide *p* ≥ 0.01 and *Viterbi p* ≥ 0.01) upon the divergence of the gloeosporioides-complex clade include GH43, GH106, CBM50 and AA7. At the divergence of the acutatum-complex clade, there were 22 expansions, of which expansions in GH78, GH43 families were rapid and significant and there was only one contraction. The divergence event of the graminicola-complex clade involved contractions in many CAZyme families with pectin degradation activity showing significant, rapid contractions (family-wide *p* ≥ 0.01 and *Viterbi p* ≥ 0.01)in families AA7, CBM50, CE8, GH28, GH78, PL1, and PL3. Divergence of the destructivum complex-clade was associated with 11 expansions and 21 contractions, of which expansion in AA7, GH74 and CE10 was significant and rapid.

Among the other species considered, *Fusarium oxysporum* had the highest number of genes (*n*=344) that were gained, with 75 expanded CAZyme with respect to its MRCA (S20 Table). *Colletotrichum incanum* had the second highest number of gene family expansions (*n*=35) and genes gained (*n*=69) followed by *C. higginsianum* (31 and 68 respectively). The highest number of contracted CAZyme families was identified in *Sordaria macrospora* (*n*=87) with a loss of 219 genes compared to the ancestral node. Forty CAZyme families contracted and only nine expanded in *C. tanaceti* with respect to the MRCA with *C. higginsianum*. The AA2 family with lignin peroxidase activity and the hemicellulose degrading GH12, GH74 families were among the expanded families, but many families with pathogenicity and plant cell wall degrading activity had contracted in *C. tanaceti*. However, the highest number of significant, rapidly evolving gene families was reported from *C. tanaceti* (*n*=9) followed by *F. oxysporum* and *C. higginsianum*, both which had seven rapidly evolving gene families each. In *C. tanaceti*, rapidly evolving CAZyme families included AA9, GH131 with lignocellulose degrading activity, chitin binding molecule families CBM18 and CBM50, GH18 with chitinase activity, GH3 and GH74 with hemicelluloses degrading activity, GH78 with pectinase activity and GT1 with glucuronosyltransferase activity. However, CBM18 and GH74 were the only families that expanded among those above with the rest contracting in *C. tanaceti* with respective to their MRCA. Gloeosporioides complex species had the largest ‘CAZyme pathogenicity profiles’ among all *Colletotrichum* species considered. The CAZyme pathogenicity profile of *C. tanaceti* was most similar to those of *Colletotrichum* species known to have an intermediate host range, infecting many hosts within a single plant family or few hosts across several plant families (Fig 4). When compared the overall pathogenicity gene profiles of all *Colletotrichum* spp., which included the numbers of the SMB clusters, transporters, P450s, CAZymes and the homologs to the PHI database, the profile of *C. tanaceti* was most similar to *C. orchidophilum* and *C. chlorophyti* (Fig 5).

**Fig 4.**
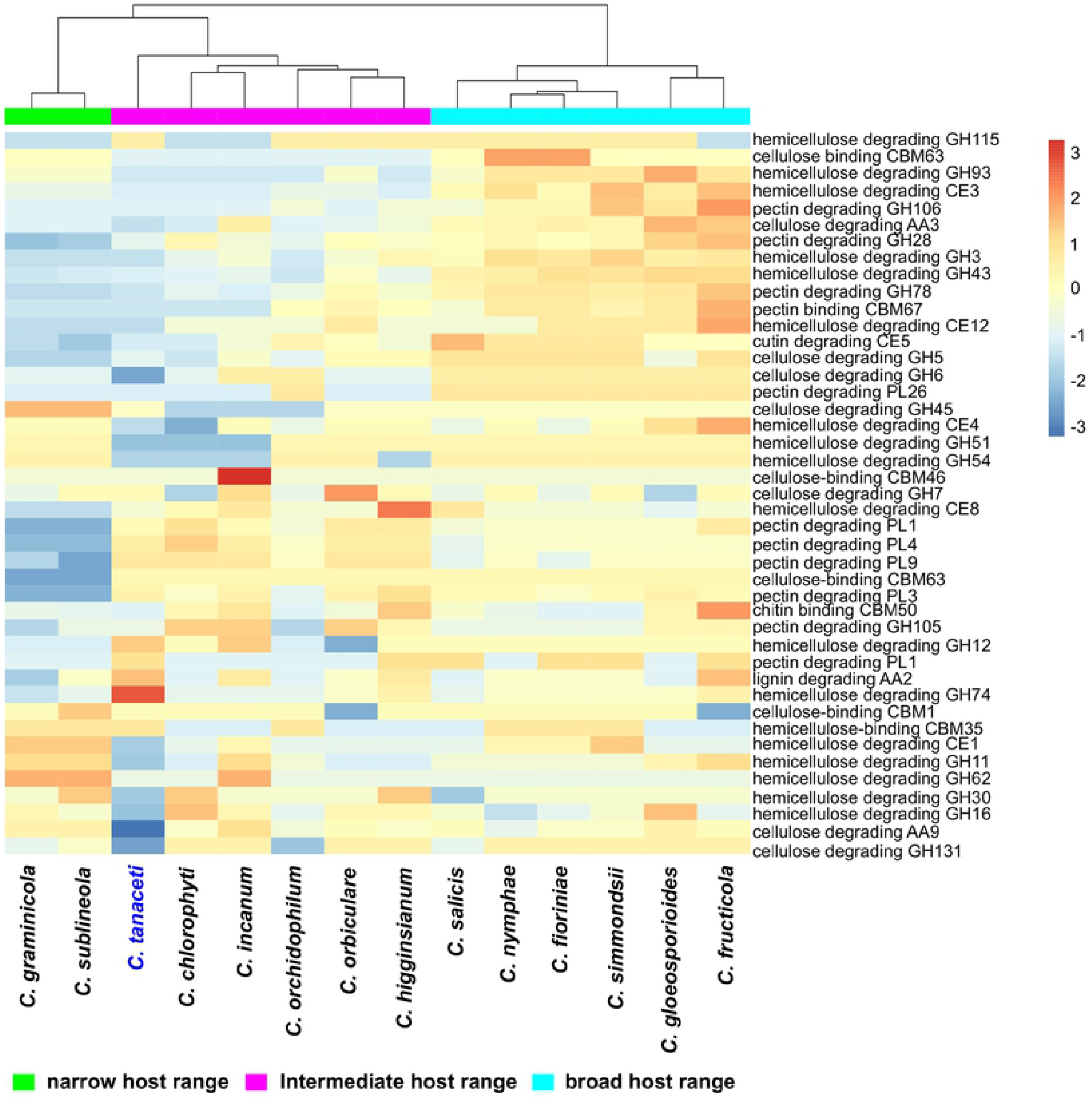
Comparison of CAZyme pathogenicity profiles predicted for *Colletotrichum* species. Hierarchical clustering performed with Euclidean distance and Ward linkage.

**Fig 5.**
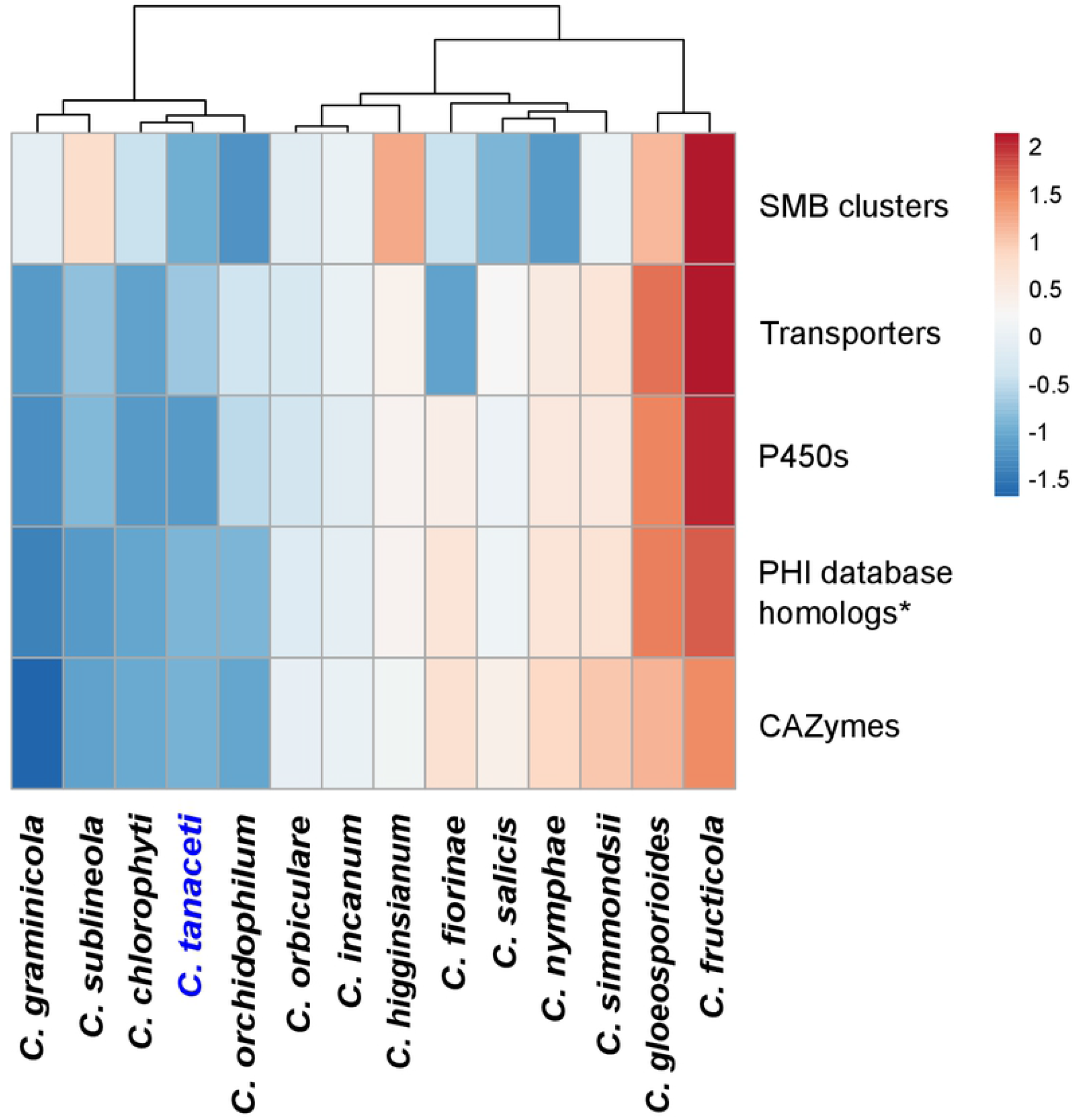
Comparison of the overall pathogenicity profiles predicted for *Colletotrichum* species. The numbers of CAZymes, secondary metabolite biosynthetic gene clusters (SMB), homologs in the transporter classification database (transporters), homologs in the fungal cytochrome P450 database (P450) and the number of homologs in the PHI database, excluding the homologs to entries in the “unaffected pathogenicity” database were used in the analysis. Hierarchical clustering was performed using Euclidean distance and Ward linkage methods.

#### Relationship of pathogenicity-related gene categories and repeat elements

The permutation tests confirmed that genes in all the tested pathogenicity-related gene categories are located significantly closer to tandem repeats than expected in a random sample (Table 4). The negative *Z*-scores confirmed the mean distance between those genes and the nearest repetitive element was less than mean of a random sample of the genome. Furthermore, all gene categories except the CAZymes were located significantly closer to the interspersed repeats. However, the expanded and the contacted subgroups of the total CAZome were significantly associated with interspersed repeats (Table 4).

**Table 4.** Permutation tests for association of repetitive elements with pathogenicity gene categories

| Gene categories of interest | All repeats <sup>a</sup> |  | Tandem repeats |  |  | Interspersed repeats |  |
| --- | --- | --- | --- | --- | --- | --- | --- |
|  | Z score <sup>b</sup> | P value <sup>c</sup> | Number of overlaps | Z score <sup>b</sup> | P value <sup>c</sup> | Z score <sup>b</sup> | P value <sup>c</sup> |
| CAZymes | -5.97 | ≤0.001 | 554 | -3.914 | ≤0.001 | -0.443 | 0.334 |
| Expanded CAZymes | -3.514 | ≤0.001 | 96 | -4.553 | ≤0.001 | -3.050 | ≤0.001 |
| Contracted CAZymes | -4.413 | ≤0.001 | 240 | -3.237 | ≤0.001 | -5.883 | ≤0.001 |
| Effectors | -5.631 | ≤0.001 | 98 | -4.725 | ≤0.001 | -3.861 | ≤0.001 |
| Peptidases | -5.787 | ≤0.001 | 82 | -4.679 | ≤0.001 | -3.895 | ≤0.001 |
| SMB clusters | -7.901 | ≤0.001 | 171 | -8.490 | ≤0.001 | -2.610 | 0.003 |

<sup>a</sup> tandem and interspersed repeats
<sup>b</sup> Z-statistic estimate and its <sup>c</sup> associated probability computed based on 10,000 random iterations.

#### Accumulation of Pathogenicity genes in the A-T rich regions of *C. tanaceti* genome

Distinct A-T rich regions and G-C equilibrated regions were identified in the genome of *C. tanaceti* (Fig 6). A total of 24.3% of the genome which had an average length of 3.77 Kb was rich in A-T and had a maximum G-C of 29%. A total of 85 genes were reported in these regions which had a gene density of 6.04 genes per Mb but the majority (68.25%) of these genes was hypothetical. Two secondary metabolite biosynthetic genes, 3 CAZymes, 2 cytochrome P450s, 2 lipases, 4 transporters, one transcription factor and one DNA polymerase were also among the genes in the A-T rich regions (S21 Table). The G-C equilibrated regions accounted for 75.7% of the genome and the average length was 14.6 Kb. The maximum G-C percentage in these regions was 55.6 and 12,087 genes were reported with a gene density of 276 genes per Mb.

**Fig 6.**
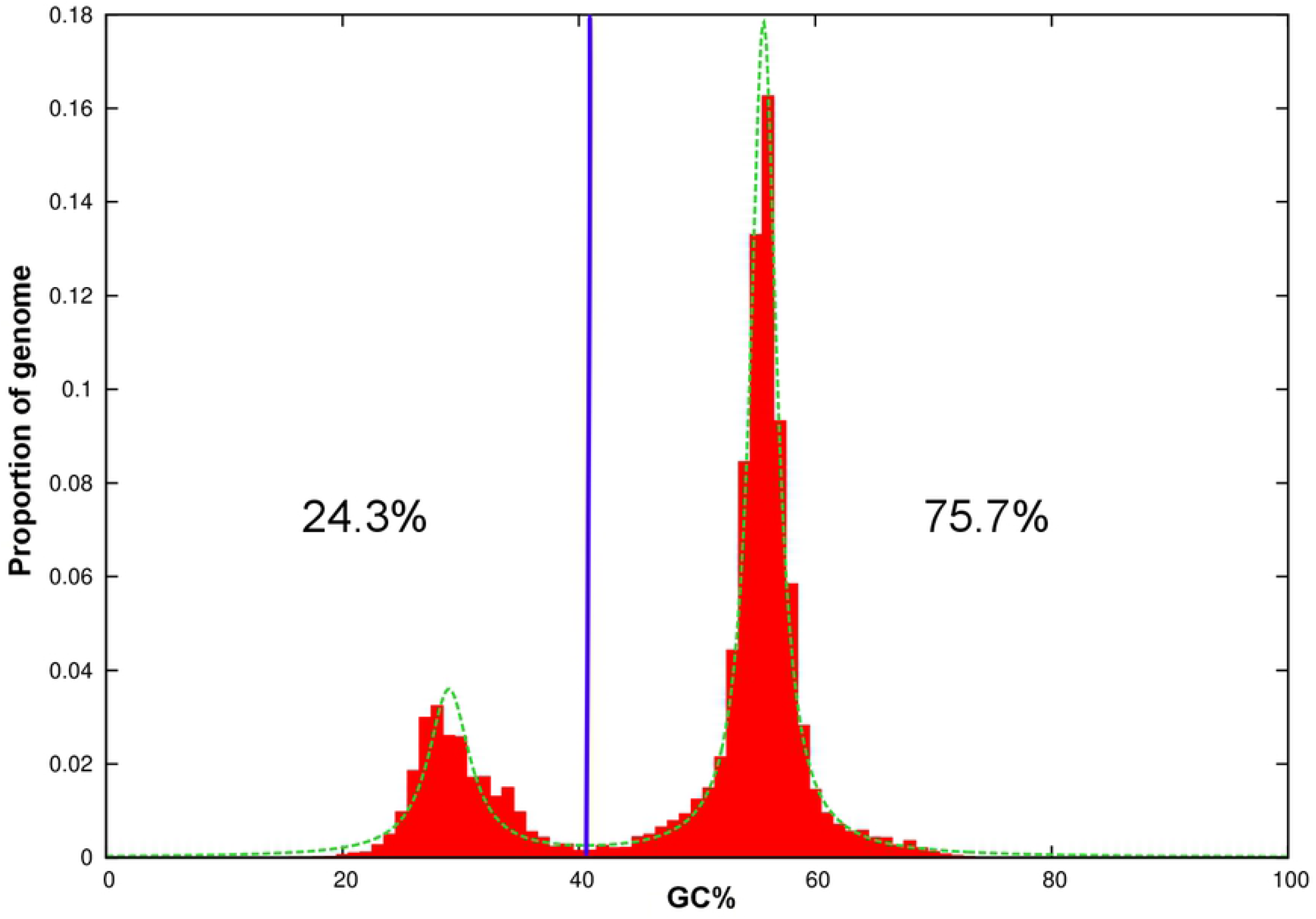
Plot of GC-content in the draft genome of *Colletotrichum tanaceti* against proportion of the genome. Genome segments were classified into A-T rich (24.3%) and G-C equilibrated (75.7%) using a GC content threshold of 40% (vertical blue line).

## DISCUSSION

### Genome and the repeat content of *Colletotrichum tanaceti*

This study reports the first draft genome sequence and annotations of the emerging plant pathogen, *C. tanaceti*. The high N50 value and BUSCO completeness indicates the high quality of the assembly and AED scores of less than one for the majority of predicted genes (93.3%) suggested that these genes had at least partial congruence with the transcriptomic evidence [114]. These good quality gene predictions and annotations will provide a solid foundation for downstream genetic, population genomic and evolutionary studies.

The genome of *C. tanaceti* had a larger repeat content (25%) than the typical 3-10% in fungi [115]. Simple sequence repeats comprised 3.03% of the genome of *C. tanaceti* which itself was unusually high for fungi (generally 0.08-0.67%) [116]. However, the majority of repeats were interspersed transposable elements (TE) (21%). TE content of *C. tanaceti* was higher than in six previously studied *Colletotrichum* species, including *C. higginsianum* which is in the same species complex, but lower than in *C. orbiculare* (44.8%). The majority of TE were retro-transposons, similar to other *Colletotrichum* spp. [117]. Proliferation of repetitive elements especially transposons, is known to be a major mechanism driving expansion of eukaryote genomes [118, 119] and therefore, could be the reason the *C. tanaceti* genome is larger than average for fungi in the phylum Ascomycota (36.91 Mb) [120].

Repeat-induced-point mutation (RIP) is a fungal-specific mechanism for limiting transposon proliferation below destructive levels [39]. RIP is known to generate A-T rich regions with lower gene densities and higher evolutionary rates than the core genome, thus generating “two-speed” genomes in several fungi [117, 121–123]. The presence of A-T rich, gene sparse regions in the *C. tanaceti* genome could therefore, be a byproduct of the RIP due to TE proliferation. Accumulating repeats followed by expanding genome size with respect to the non-pathogenic strains is a trend observed in many plant pathogenic fungi and can provide an evolutionary advantage in terms of pathogenesis [124]. The high repeat content of *C. tanaceti* may have an important role in generating genome plasticity [125].

### Pathogenicity genes of *C. tanaceti*

A large array of putative genes related to pathogenicity was inferred from the sequenced genome of *C. tanaceti*. Apart from many plant cell wall-degrading enzymes, effectors, *P450s* and the proteolytic enzymes, there were proteins with CFEM domain (pfam05730) [126] with roles in conidial production and stress tolerance [127] among the secreted proteins. The average cysteine composition, length and proportion of specificity of the candidate secreted effectors of *C. tanaceti* were similar to those hemibiotrophic pathogens [128]. However, a minority of effector candidates was neither small (<300bp) nor rich in cysteine (>3%), similar to previous reports of atypical effectors [129]. Effector candidates with a nuclear localization signal might translocate to the host nucleus and reprogram the transcription of genes related to host immune responses. Homologs to known effectors, and effectors with conserved domains of virulence factors may have similar functions in *C. tanaceti*, for example, in penetration peg formation (cyclophllin) [130], phytotoxity induction (cerato-platanin) [131] and adherence of the fungal structures to other organisms (hydrophobin) [132].

Most secreted proteases of *C. tanaceti* were serine proteases predicted to evade plant immune responses by degrading plant chitinases [22]. Subtilisins (S08) were the most abundant of these in *C. tanaceti*, similar to reports in other fungi [22]. Subtilisins, with their alkaline optima, and the proteases in other subfamilies with acidic optima, such as A01, C13, G01, M20 and S10 [133], might enable *C. tanaceti* to degrade plant proteins across a wide pH range. Also, the protease inhibitors of *C. tanaceti* might have effector-like roles via inhibition of plant defense proteases [134].

The SMB gene clusters and the candidate proteins of MAPKs pathways identified in the genome of *C. tanaceti* are also believed to play an important role in pathogenesis. The majority of the secondary metabolite clusters of *C. tanaceti* were type 1 PKs-like which are usually associated with synthesizing fungal toxins. Melanin, another important secondary metabolite aids penetration via increasing turgor pressure [135]. Even though the gene cluster associated with melanin biosynthesis was not identified, the homolog of the melanin biosynthetic gene *SCD1* encoding Scytalone dehydratase [136] in *C. tanaceti* is worth investigating further since *SCD1* has been successfully used as a target for fungicides to control other pathogens [137]. Apart from their function in SM biosynthesis, the candidate P450s of *C. tanaceti* could be involved in housekeeping roles and therefore, could be good targets for fungicide development, as in the case of azoles targeting CYP51 [138]. Furthermore, the candidate proteins of MAPKs pathway in *C. tanaceti* could play a crucial role in appressorium formation [25, 139], penetration [140], conidiation [141] and pathogenesis-related morphogenesis [142], as reported for *C. higginsianum* and *C. lagenaria*.

Of the CAZyme families identified to be expanded in *C. tanaceti*, the chitin binding family CBM18 could play a role in protecting the *C. tanaceti* cell wall from exogenous chitinases, as is the case in *Trichoderma reesei* [143]. The expansion of the hemicellulose-degrading GH74 family could promote rapid degradation of host tissues by *C. tanaceti* during the necrotrophic phase. The expansion of the lignin-degrading AA2 family in *C. tanaceti* has the potential to assist infection of xylem vessels and thereby aid translocation of propagules to different parts of the plant and establishing secondary infections.

The conserved nature of certain pathogenicity genes, such as the secondary metabolite clusters within the destructivum complex, was evident with their presence within the syntenic blocks with *C. higginsianum*. However, only a minority of the effectors, proteases and SM backbone genes of *C. tanaceti* were among the core gene set for *Colletotrichum* overall, therefore emphasizing their role in adaptation to new hosts. The species-specific effectors, singletons from the orthology analysis and the genes exclusive to *C. tanaceti* might have been horizontally transferred or be related to the host affiliation and niche specialization of *C. tanaceti*. Taken together, this inferred pathogenicity gene suite of *C. tanaceti* could be targeted in future resistance breeding and other disease management strategies for *C. tanaceti*.

### Host range of *Colletotrichum tanaceti*

The proposed pathogenicity gene repertoire of *C. tanaceti* was most similar to that of pathogens with intermediate host ranges. The number of pathogenicity genes inferred from *C. tanaceti* was either similar to or less than the average for all *Colletotrichum spp*. investigated but the overall composition was similar to *Colletotrichum* spp. which either were able to infect many species within a plant family or few species across families. The comparison of CAZyme pathogenicity profiles among *Colletotrichum* spp., with both expansions and contractions with respective to its MRCA clearly suggested an intermediate host range for *C. tanaceti*.

The pathogenicity profile of *C. tanaceti* was very distinct from that of the other destructivum complex member, *C. higginsianum*, despite the two species sharing the highest number of orthologs and having the shortest evolutionary distance. Contractions in many pathogenicity gene families in *C. tanaceti* compared to *C. higgginsianum* indicated more restricted pathogenicity in *C. tanaceti*. The most similar CAZyme pathogenicity profile to that of *C. tanaceti* was from *C. chlorophyti* which has been reported to infect herbaceous hosts such as tomato (plant family Solanaceae) and soybean (plant family Fabaceae) [69]. The similarity to *C. chlorophyti* was consistent for other gene categories such as the P450s, transporters and the overall pathogenicity profile. A homolog to the demethylase (PDA), which provides tolerance to the phytoalexin pisatin synthesised by *Pisum sativum* [144], was inferred in *C. tanaceti* (CTA1_6324s) which could be an indicator of the ability of *C. tanaceti* to infect Fabaceae. The composition of the SMB cluster was however, more similar to *C. orchidophilum*, another pathogen reported to infect the herbaceous, monocot plant family of Orchidaceae [145]. The similarity of the pathogenicity profile of *C. tanaceti* to two pathogens infecting multiple herbaceous plant species was notable as the only known host of *C. tanaceti* is also herbaceous. Both *C. chlorophyti* and *C. orchidophilum* have been reported from multiple host species. Therefore, the pathogenicity gene suite of *C. tanaceti* suggests that *C. tanaceti* has the genetic ability to infect more hosts than currently recognized. If *C. tanaceti* can infect other hosts, such crops could also provide an external gene pool of inoculum for infection of pyrethrum crops increasing the evolutionary potential of the pathogen populations. Based on results of comparative analysis of pathogenicity profiles, a further hypothesis is that these alternative hosts are likely to be herbaceous plants. Future studies investigating the cross-host infectivity and pathogenicity of *C. tanaceti* are recommended.

### Evolution of pathogenicity genes

Pathogenicity genes of *C. tanaceti* appear to be capable of evolving relatively rapidly. Tandem repeats such as simple sequence repeats have high mutation rates [146] and could promote frameshift mutations in adjacent genes by slipped misalignment during replication. Therefore, the significant overlap between the tandem repeats and the pathogenicity genes suggested high potential to mutate and create different pathotypes. Transposons promote insertional mutations that can either cause disruption or modification of gene expression or generate new proteins and also are major drivers of gene duplication [147]. Transposons were in close proximity to gene categories of pathogenicity in *C. tanaceti* such as the SMB clusters, peptidases and effectors. The significant association of TE with pathogenicity genes were previously reported in *C. truncatum* [117] and *C. higginsianum* [15].

*Colletotrichum tanaceti* had the highest number of rapidly evolving CAZyme families among the 17 species studied which also was indicative of the rapid evolutionary rate in these pathogenicity genes. Interspersed repeats were not in close proximity to the total CAZome. They were however, located significantly closer to the expanded or contacted families indicating that interspersed repeats were a major contributor to CAZyme family expansions/contractions in *C. tanaceti* by causing gene duplication (in expansions) or gene disruptions (in contractions) [118, 148]. Although gene sparse, the A-T rich regions of the genome contained several (*n*=18) known pathogenicity and virulence factors and many hypothetical proteins which could be facilitating adaptive evolution. According to the two-gene hypothesis the genes in the A-T rich regions can evolve faster than the ‘core’ genome [124]. Duplication of pathogenicity and virulence genes and a higher mutation rate may allow more rapid pathogen responses to evolution of resistance in existing hosts or adaptation to new host species.

### Genus *Colletotrichum*

Phylogenetic relationship throughout the genus was consistent with previous observations, with gloeosporioides complex members and *C. orbiculare* forming a clade separately from the destructivum, graminicola and acutatum clades [9–11]. One notable difference was in the divergence time estimates for the divergence of *Colletotrichum* species complexes which were more ancient than reported by Liang et al [11], despite using the same calibration times. This could have been due to this study using cross-validation across 50 smoothing factors in CAFÉ as opposed to using 12 different constraints and smoothing factor combinations differences, as the use of the different data sets.

Comparative genomic analyses emphasized the rapid evolutionary rate and the high diversity within the genus. The short time for speciation within the acutatum complex, and the fourteen *Colletotrichum* species in general, was suggestive of the high evolutionary rate within the genus with respective to the typical evolutionary rate of the fungal kingdom (0.0085 species units per Myr) [149]. The sequence similarity between *C. tanaceti* and other species of *Colletotrichum* varied widely and dropped drastically with evolutionary distance, suggesting high diversity within the genus. However, the drop in orthology was less dramatic, emphasizing the contribution of non-coding regions in generating diversity within the genus. The extent of synteny between *C. tanaceti* and *C. higginsianum* was high and very similar to the percentage synteny previously reported for the two graminicola complex species, *C. sublineola* and *C. graminicola* [150]. This suggested that even though there was high diversity within the genus, the species in the same species complex tend to share more synteny than the between species complexes.

Evolutionary analysis of CAZyme families of different *Colletotrichum* lineages revealed an association between CAZyme families and host range. The GH131 with cellulose degrading activity was the only rapidly evolving gene family at the MRCA of *Colletotrichum* spp. suggesting a possible association of this family with speciation and host determination within the genus. Families GH43, with hemicellulose degrading activity and AA7, with gluco-oligosaccharide activity significantly expanded upon divergence of both the gloeosporioides and acutatum-complex clades, which could have broadened the host ranges of members of these two complexes. The significant expansions in pectin degrading enzyme families GH106 in gloeosporioides and GH78 in the acutatum clades could also have enabled degradation of pectin rich cell walls of young fruits [151] of these fruit-rotting species.

The most significant contractions were reported in pectin degrading families upon the divergence of the graminicola complex clade. This could have been the reason for species in this complex exclusively infecting monocot plant species considering that the pectin content of monocot cell walls is generally less than in dicots [152]. Even though this was a similar result to previous studies [7], *C. orchidophilum* which is known to infect plants from monocot family Orchidaceae [153], deviated from this pattern. Gene family AA7 was rapidly evolving in many *Colletotrichum* species and could have been involved in biotransformation or detoxification of the lignocellulosic compounds [154].

In general, the overall CAZyme pathogenicity profiles of *Colletotrichum* spp. followed host range of those species rather than the taxonomy. The gloeoporioides and acutaum complex members which have broad host ranges, but are evolutionary distant, were clustered together. This could be a byproduct of the “two-speed” genome scenario in certain *Colletotrichum* spp. such as *C. orbiculare, C. chlorophyti, C. graminicola* [117] and as suggested, also in *C. tanaceti*. In this scenario, the pathogenicity genes are located in repeat-rich regions, allowing them to evolve at a higher rate than the rest of the genome. This was also evident by the significant association of TE with pathogenicity genes in *C. tanaceti* and in *C. truncatum* [117] and *C. higginsianum* [15]. This scenario would cause the species with similar pathogenicity gene profiles to cluster together, despite their evolutionary distance.

## CONCLUSION

In conclusion, a draft genome of *C. tanaceti* was used to quantify the molecular basis of pathogenicity of the species and to improve the knowledge of the evolution of the fungal genus *Colletotrichum. Colletotrichum tanaceti* is likely to have alternative hosts and is a potential threat to the crops grown in rotation with pyrethrum. The genome of *Colletotrichum tanaceti* contains a large component of repetitive elements that may result in genome expansion and rapid generation of novel genotypes. The tendency of the pathogenicity genes to evolve rapidly was evident in genomic signals of the RIP and association of repeats with the pathogenicity genes. Therefore, with a large array of pathogenicity genes that potentially can evolve rapidly, *C. tanaceti* is likely to become a high-risk pathogen to global pyrethrum production. Complexity of the *Colletotrichum* genus was evident with its high diversity and evolutionary rate. The significant expansions and contractions of gene families upon divergence of different lineages within the genus could be important determinants in species complex diversification in *Colletotrichum*. The reason for pathogenicity genes to have different clustering than the phylogeny in *Colletotrichum* could be the occurrence of “two-speed” genomes in certain species. These findings will facilitate future research in genomics and disease management of *Colletotrichum*.

## ACKNOWLEDGEMENTS

We thank Dr Kym Pham and Dr Arthur Hsu for performing library preparation, sequencing and genome assembly which were conducted at the Melbourne Translational Genomics Platform (Department of Pathology, University of Melbourne). We also thank Botanical Resources Australia - Agricultural Services, Pty. Ltd for supporting the project. RVL received Melbourne International Fee Remission Scholarship and Melbourne International Research Scholarship from the University of Melbourne, Australia. NDY was supported by a Career Development Fellowship (CDF) from NHMRC. PKK was supported by an Early Career Fellowship (CDF) from NHMRC.

## Supplementary information

**S1 Table.** GO term enrichment analysis in *C. tanaceti*

**S2 Table.** KEGG orthology annotations of *C. tanaceti*

**S3 Table.** KEGG pathway map IDs of *C. tanaceti*

**S4 Table.** KEGG orthology assignments of Map kinase pathway in *C. tanaceti*

**S5 Table.** KEGG orthology assignments of Aflatoxin biosynthesis pathway in *C. tanaceti*

**S6 Table.** KEGG orthology assignments of ABC transporters in *C. tanaceti*

**S7 Table.** Global alignment of *C. tanaceti* contigs to the *C. higginsianum* chromosomes

**S8 Table.** Secreted proteins of *C. tanaceti* and their conserved domains

**S9 Table.** Secreted effector candidates of *C. tanaceti* with homology to known effectors and conserved motifs

**S10 Table.** Secreted peptidases of *C. tanaceti*

**S11 Table.** Secreted peptidase inhibitors of *C. tanaceti*

**S12 Table.** Secondary metabolite bioysnthetic gene cluster predtions of *C. tanaceti*

**S13 Table.** Conserved domains of secondary metabolite biosynthetic genes of *C. tanaceti*

**S14 Table.** Homologs in *C. tanaceti* to fungal cytochrome P450 database

**S15 Table.** Homologs in *C. tanaceti* to transporter classification database

**S16 Table.** Homologs to pathogen host interaction database in *C. tanaceti*

**S17 Tabel.** CAZyme family assignment of *C. tanaceti*

**S18 Table.** Family-wide probability values and viterbi probability values of CAZyme families across taxa

**S19 Table.** Expansions and contractions of CAZyme familes upon divergence of different lineages

**S20 Table.** Statistics of CAZyme gene family evolution across taxa

**S21 Table.** Genes in the Atrich region of *C. tanaceti* genome

**S1 Fig. Median GC percentage and median total length (Mb) (x axis) of publicly available draft genomes representing *Colletotrichum* species (y axis).**

**S2 Fig. Circular plot showing synteny between *Colletotrichum tanaceti* contigs (numbers) mapped to the 12 individual chromosomes (NC_codes) of the *C. higginsianum* genome.**

**S3 Fig. Comparison of type of secondary metabolite biosynthetic gene clusters** (gene clusters producing Terpenes, Indoles, Polyketides (PKs), Non-ribosomal peptides (NRPs_and hybrids of the above categories) in *Colletotrichum* species; hierarchical clustering performed using Euclidean distance and Ward linkage.

**S4 Fig. Comparison of composition of pathogen host interaction database (PHIbase) homolog profiles** (number homologs to entries in “reduced virulence”, “unaffected pathogenicity”, loss of pathogenicity”, “effector”, “lethal” and “increased virulence” categories in the PHIbase) in *Colletotrichum* and related species; hierarchical clustering performed with Euclidean distance and Ward linkage.

